# Towards a fully automated surveillance of well-being status in laboratory mice using deep learning

**DOI:** 10.1101/582817

**Authors:** Niek Andresen, Manuel Wöllhaf, Katharina Hohlbaum, Lars Lewejohann, Olaf Hellwich, Christa Thöne-Reineke, Vitaly Belik

## Abstract

Assessing the well-being of an animal is hindered by the limitations of efficient communication between humans and animals. Instead of direct communication, a variety of behavioral, biochemical, physiological, and physical parameters are employed to evaluate the well-being of an animal. Especially in the field of biomedical research, scientifically sound tools to assess pain, suffering, and distress for experimental animals are highly demanded due to ethical and legal reasons. For mice, the most commonly used laboratory animals, a valuable tool is the Mouse Grimace Scale (MGS), a coding system for facial expressions of pain in mice which has been shown to be accurate and reliable. Currently, MGS scoring is very time and effort consuming as it is manually performed by humans being thoroughly trained in using this method. Therefore, we aim to develop a fully automated system for the surveillance of well-being in mice. Our work introduces a semi-automated pipeline as a first step towards this goal. We use and provide a new data set of images of black-furred laboratory mice that were moving freely, thus the images contain natural variation with regard to perspective and background. The analysis of this data set is therefore more challenging but reflects realistic conditions as it would be obtainable without human intervention. Images were obtained after anesthesia (with isoflurane or ketamine/xylazine combination) and surgery (castration). We deploy two pre-trained state of the art deep convolutional neural network (CNN) architectures (ResNet50 and InceptionV3) and compare to a third CNN architecture without pre-training. Depending on the particular treatment, we achieve an accuracy of up to 99% for binary “pain”/”no-pain” classification.

**Author summary:** In the field of animal research, it is crucial to assess the well-being of an animal. For mice, the most commonly used laboratory animals, there is a variety of indicators for well-being. Especially the facial expression of a mouse can give us important information on its well-being state. However, currently the surveillance of well-being can only be ensured if a human is present. Therefore, we developed a first approach towards a fully automated surveillance of the well-being status of a mouse. We trained neural networks on face images of black-furred mice, which were either untreated or underwent anesthesia or surgery, to distinguish between an impaired and unimpaired well-being state. Our systems successfully learnt to assess whether the well-being of a mouse was impaired and, depending on the particular treatment, its decision was correct in up to 99%. A tool that visualizes the features used for the decision making process indicated that the decision was mainly based on the facial expressions of a mouse.

## Introduction

The Directive 2010/63/EU stipulates to fully apply the 3-R-principle of Russel and Burch [1] with regard to animal experimentation. Wherever possible animal experiments shall be replaced by alternative methods. However, if an animal experiment is deemed necessary, the total number of animals used in the experiment is to be reduced to a minimum necessary to obtain scientifically sound results. Moreover, refinement measures shall be applied to foster the well-being of the laboratory animals. Refinement measures aim to minimize pain, suffering, and distress accompanying the experiment. Moreover, refinement significantly contributes to the quality of research findings [2]. Therefore, in the scope of animal welfare and good science, it is crucial to develop scientifically sound tools which help to systematically evaluate and improve the well-being of laboratory animals.

Impaired well-being of animals can be assessed by behavioral, biochemical, physiological, and physical parameters [3]. In recent years, methods to analyze the facial expressions of pain, analogous to the facial action coding system (FACS) for humans [4], were developed for various animal species, e.g., mice, rats, rabbits, cats, horses, cows, sheep, and piglets [5–14]. The so-called Grimace Scales are thought to measure the presence or absence of grimacing associated with pain. Especially for mice, the most commonly used laboratory animals, the Grimace Scale became a valuable tool and was applied in pain research as well as cage-side clinical assessment [15–17]. However, prerequisite to apply the Mouse Grimace Scale (MGS) is that a person is present and either generates live scores or acquires images or videos to be scored retrospectively. Thus, during periods in which the animals are not monitored, the well-being of a mouse cannot be assessed. Another important aspect is the fact that mice are prey animals and often hide signs of weakness, injury, and pain in the presence of humans [18]. Therefore, it is decisive to find a way to automatically monitor well-being of a mouse in the absence of humans. Since the facial expression has proven to be useful in mice, it is worthwhile to develop an automated facial expression recognition software. Automation can also bring substantial time savings for the MGS application as it is extremely labor intensive to train persons in using the MGS and to manually generate MGS scores.

In the field of automatic facial expression recognition in animals, we are only at the very beginning. In contrast, a lot of advances have already be made in automated facial expression recognition in humans [19]. Recently, machine learning techniques experienced a revival, mostly due to the progress in deep learning (multi-layered neural networks combined with advanced optimization techniques) [20]. They allow to perform classification and predictions on the data without *a priori* feature design. Statistical physics recently provided evidence for their unexpected effectivity [21]. Deep learning is used in different fields of research, relating to image recognition, from urban dynamics [22, 23] to tracking of humans and animals in general [24], which is important in neuroscience, behavioral biology and digital pathology [25]. These networks are successfully used in many applications involving the acquisition of information about features of an image that traditionally only humans or animals could perceive. Two influential examples are AlexNet [26] and R-CNN [27]. A number of highly efficient computational frameworks for deep learning and neural networks were recently released, e.g., TensorFlow, Theano, and Caffe. They are easy to deploy and make the technology accessible for a vast audience of researchers [28]. Furthermore, these libraries are equipped with a range of pre-trained models allowing the efficient learning of high level concepts using transfer learning strategies.

The goal of our study was to develop an automated facial expression recognition software for mice that is able to assess whether the well-being of a mouse is impaired by post-anesthetic or post-surgical pain and/or distress. Therefore, we use a pipeline with two components. First, a detector that locates and extracts a face of a mouse in the processed image and secondly, a deep neural network model for the classification of facial expressions. We used an own image data set of adult female and male black-furred mice of the strain C57BL/6JRj which were either untreated or received anesthesia (with isoflurane or ketamine/xylazine combination) with or without surgery (castration) [29, 30].

## Materials and methods

### Ethics statement

In the present study, we reused images from mice which were obtained in previous animal experiments [29, 30]. Animal experimentation was performed according to the guidelines of the German Animal Welfare Act and the Directive 2010/63/EU for the protection of animals used for scientific purposes. Maintenance of mice and all animal experimentation were approved by the Berlin State Authority (“Landesamt für Gesundheit und Soziales”, permit number: G0053/15).

### Animals

Images of mice were obtained from previous studies performed at Freie Universität Berlin (Berlin, Germany) using 61 female and 65 male adult C57BL/6JRj mice purchased from Janvier Labs (Saint-Berthevin Cedex, France) [29, 30]. At the age of 10–13 weeks mice either underwent inhalation anesthesia with isoflurane (26 male and 26 female C57BL/6JRj mice) [29], injection anesthesia with the combination of ketamine and xylazine (26 male and 22 female C57BL/6JRj mice) [30], or no treatment (13 male and 13 female C57BL/6JRj mice). Additionally, 19 male C57BL/6JRj mice at the age of 18–42 weeks, which had previously received injection anesthesia or no treatment, were castrated at the age of 18–42 weeks in order to be re-socialized in groups of 3–4 animals.

Female mice were group-housed with 3–5 mice in Makrolon type IV cages (55 × 33 × 20 cm). Male mice had to be single-housed in Makrolon type III cages (42 × 26 × 15 cm) due to aggressive behavior toward conspecifics. The cages contained fine wooden bedding material (LIGNOCEL® 3–4 S, J. Rettenmaier & Söhne GmbH + Co. KG, Rosenberg, Germany) and nest material (nestlets: Ancare, UK agents, Lillico, United Kingdom; additionally, cocoons were provided for castrated mice: ZOONLAB GmbH, Castrop-Rauxel, Germany). A red plastic house (length: 100 mm, width: 90 mm, height: 55 mm; ZOONLAB GmbH, Castrop-Rauxel, Germany) and metal tunnels (length: 125 mm, diameter: 50 mm; one tunnel in Makrolon type III cages, two tunnels in Makrolon type IV cages) were provided as cage enrichment. The animals were maintained under standard conditions (room temperature: 22 ± 2 ° C; relative humidity: 55 ± 10%) on a light:dark cycle of 12:12 h of artificial light with a 5 min twilight transition phase (lights on from 6:00 a.m. to 6:00 p.m.). The mice were fed pelleted mouse diet ad libitum (Ssniff rat/mouse maintenance, Spezialdiäten GmbH, Soest, Germany) and had free access to tap water. Both the technician and veterinarian were female. Combined tunnel and cup handling were used, i.e., the mice were carefully caught in a tunnel and then transferred to the hand, in order to minimize handling induced stress and anxiety [31]. After the study, females as well as castrated males were re-homed and intact male mice were used for educational purposes.

### Animal experimental procedures

#### Inhalation anesthesia with isoflurane

Inhalation anesthesia was induced with 4% isoflurane (Isofluran CP®, CP-Pharma Handelsgesellschaft mbH, Burgdorf, Germany) in 100% oxygen in an anesthetic chamber (with sliding cover, Evonik Plexiglas, 240 × 140 × 120 mm). The chamber was not prefilled with isoflurane. After the mouse had lost the righting reflex, it was transferred to a heating pad and anesthesia was maintained with 1.75–2.5% isoflurane in 100% oxygen via nose cone. Artificial tears (Artelac® Splash MDO®, Bausch & Lomb GmbH, Berlin, Germany) were administered to both eyes to prevent the eyes from drying out. Anesthesia lasted for approximately 45 minutes. During anesthesia pedal withdrawal and lid reflex were regularly tested and vital parameters (i.e., respiratory rate, heart rate, and oxygen saturation) were monitored. According to the design of our previous study, mice were anesthetized either once or six times at an interval of three to four days [29].

#### Injection anesthesia with the combination of ketamine and xylazine

For injection anesthesia a stock solution with 160 *µ*L Ketavet® 100 mg/mL (Zoetis Deutschland GmbH, Berlin, Germany), 160 *µ*L Rompun® 2% (Bayer Vital GmbH, Leverkusen, Germany), and 1680 *µ*L physiologic saline solution was prepared in a syringe. A dosage of 80 mg/kg ketamine and 16 mg/kg xylazine [32], warmed to body temperature, was administered intraperitoneally at a volume of 10 *µ*L/g body weight using 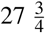 Gauge needles. Then the mouse was transferred to a Makrolon Typ III cage placed on a heating pad. After the loss of righting reflex, the mouse was transferred to a heating pad. Artificial tears (Artelac® Splash MDO®, Bausch & Lomb GmbH, Berlin, Germany) were administered to both eyes to protect them from drying out. Overall duration of anesthesia was 60–84 minutes. Pedal withdrawal and lid reflex were regularly tested and vital parameters (i.e., respiratory rate, heart rate, and oxygen saturation) were monitored during anesthesia. According to the design of our previous study, mice were anesthetized either once or six times at an interval of three to four days [30].

#### Castration

Mice were anesthetized with isoflurane in 100% oxygen (induction: 4% isoflurane in an induction chamber, maintenance: 1.5–2.5% isoflurane via nose cone). After anesthesia induction, meloxicam (1 mg/kg body weight, s.c.; Metacam 2 mg/ml Injektionslösung für Katzen, Boehringer Ingelheim Vetmedica GmbH, Ingelheim, Germany) was subcutaneously administered and lidocaine/prilocaine ointment (Emla Creme, AstraZeneca GmbH, Wedel, Germany: 1 g contained 25 mg lidocaine and 25 mg prilocaine) was applied to the scrotum. Surgical castration was performed in supine position according to Behrens et al. [33]. The testicles were removed one by one. In brief, testicles were pushed down into the scrotal sacs and an 1 cm incision was made thorough the skin at a right angle to the midline of the scrotal sac. After a testicle was pushed out, the *vas deferens* with the blood vessels running along it was ligated with absorbable suture (3–0) and the testicle was removed. When both testicles were removed, the skin was stitched with a single button suture.

#### No treatment

Mice of the control groups in our previous studies received neither anesthesia nor surgery (i.e., no treatment). Since well-being of these mice was not affected by any medical procedure, they were expected not to display any signs of pain and/or distress.

### Dataset

The present study is based on a large data set of images of C57BL/6JRj mice. Images were obtained in previous studies, in which the impact of procedures frequently performed in animal experimentation on the well-being and stress levels of mice was systematically assessed by using the MGS, among other animal-based parameters [29, 30]. According to the treatment of the mice, our data set was divided into the three subsets: KXN (ketamine/xylazine anesthesia), IN (inhalation anesthesia with isoflurane), and C (castration).

#### Image acquisition

Images were generated as described previously in Hohlbaum et al. [34]. All images were taken in observation cages (22 × 29 × 39 cm) (Fig 1) with three white walls to contrast the black mice and one clear wall. Cages were custom-made in our facility and visually varied slightly, e.g., walls were attached to each other with dark-or light-colored material. The bottom of the cage was covered with approximately 0.5 cm bedding material and soiled bedding was scattered on top in order to minimize stress caused by the novel environment. Food pellets normally supplied as diet and a water bowl were provided.

**Fig 1.**
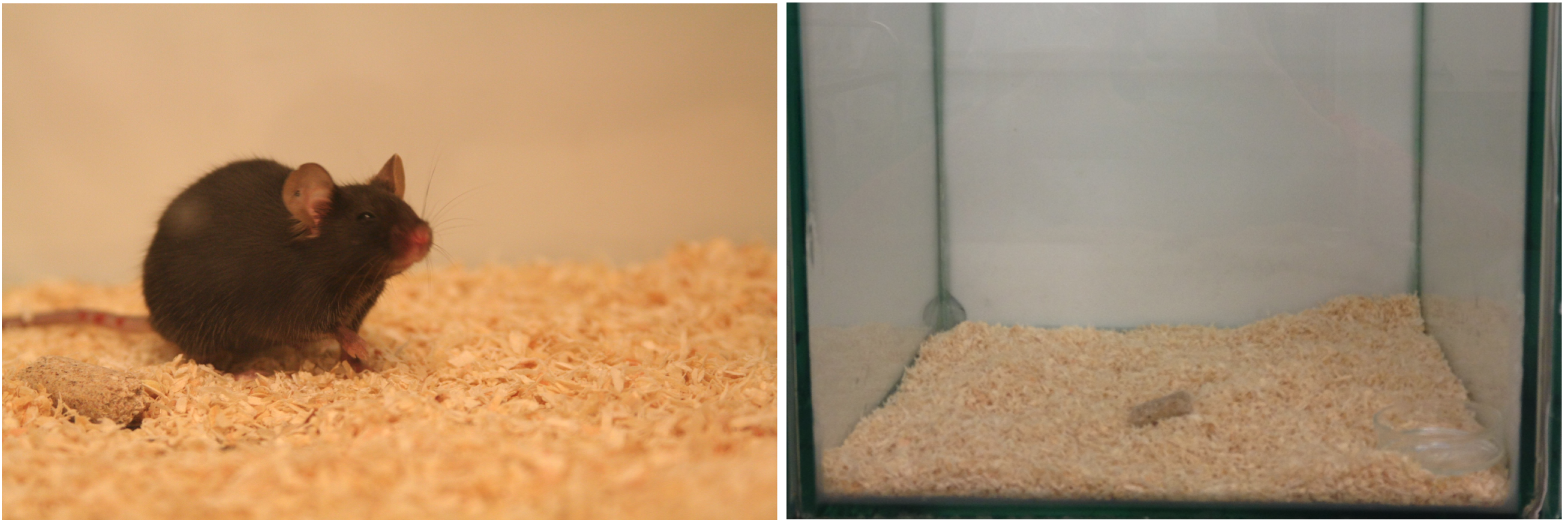
Image of the data set and observation cage. Example image of a black-furred laboratory mouse (C57BL/6JRj strain) of the dataset (left). Observation cage used for monitoring the mice after the procedures (right). Images of the mice were taken when mice were moving freely around in the observation cage.

After the procedures were performed, the mouse was gently transferred into an observation cage and was allowed to habituate to the new environment for 30 min. Then a series of images was taken within a few minutes (approximately 1-2 minutes, but in some cases even longer) using a high definition camera (Canon EOS 350D, Canon Inc., Tokyo, Japan). Baseline images were acquired prior to the procedures. Post-procedure images were taken at various points in time as follows:

- Inhalation anesthesia with isoflurane [29]: 30 min, 150 min post-procedure
- Injection anesthesia with the combination of ketamine and xylazine [30]: 30 min, 150 min, day 2, day 9 post-procedure
- Castration: 30 min, 150 min, 300 min, day 2, day 3, day 7 post-procedure.

Images of untreated mice were generated at the same times as images of corresponding treated groups. Since they were considered not to show any post-anesthetic or post-surgical distress, they were added to baseline images in the present study.

#### Human evaluation of mouse images

In order to systematically evaluate the levels of post-anesthetic or post-surgical distress and/or pain, mouse images were scored by humans on the MGS. The MGS developed by Langford et al. [5] measures characteristic changes in facial expressions of pain in mice. The five facial action units of the MGS (i.e., orbital tightening, nose bulge, cheek bulge, ear position, and whisker change) are scored on a 3-point-scale (0 = not present, 1 = moderately visible, 2 = obviously present) with high scores reflecting high intensity of a facial action unit (for further details see Langford et al. [5]). An accuracy of 72–97% for humans scorers was reported [5]. Interestingly, the change of facial action units described in the MGS is also triggered by other stimuli than pain such as post-anesthetic distress, situations associated with fear (i.e., whisker contact, social proximity, cat odor exposure, rat exposure), and aggression or subordination [29, 30, 35].

For MGS scoring, one image of high quality showing the mouse face from frontal or lateral view was randomly selected per mouse, and point in time [29, 30, 34] and the face of the mouse were manually cropped from the image so that, if possible, the body posture was not visible. Overall, five persons (three veterinarians, laboratory assistant, and secretary) with different experience levels in laboratory animal science were trained in scoring images for MGS. A minimum of two and a maximum of four individuals (IN: three, KXN: four, C: two) were recruited for MGS scoring. Images were presented in a randomized order on a computer screen to the uninformed individuals, who independently scored the five facial action units of the MGS using the manual provided by Langford et al. [5].

#### Binary “Pain” Labels

Due to the time and labor intensive process of MGS scoring, only a small amount of images (658 images) in our data set is scored on the MGS. This number does not allow a successful deep learning based regression analysis of the function, which maps from image (pixel) space to MGS score. As we could not predict the MGS score directly, we followed the approach of Tuttle et al. [36] and assigned one of two defined states, “pain” and “no pain”, to all images in order to train a binary classifier on the whole data set. To do so, time points of image acquisition were either defined as “pain” or “no pain” based on the MGS scores obtained by humans and the statistical analysis performed in our previous studies [29, 30, 34]: if MGS scores were significantly higher when compared to untreated mice (Fig 2), all images of this point in time were considered to display a “pain face” and were assigned to the label “pain” (Table 1). Furthermore, 300 min post-castration was also defined as “pain” state, though significance versus baseline was not reached. Images taken at other points in time as well as images of untreated mice were assigned to the label “no pain”. Individual MGS scores could not be considered, i.e., an image may have been labeled with “pain” even though the image shows a mouse with a low MGS score. Important to note here is that the terms “pain” and “no pain” were used as labels and do not necessarily correspond to the actual state of a mouse. This should especially be considered in the case of images generated after anesthesia. Mice which received anesthesia only and did not undergo surgery may have experienced post-anesthetic distress rather than pain [29, 30].

**Table 1.** Points in time labelled with “pain” are listed for each procedure. Baseline images and images acquired at a later time were assigned to the label “no pain”.

| Procedure | Time points assigned the label "pain" |
| --- | --- |
| Isoflurane anesthesia | 30 min post-procedure |
| Ketamine/xylazine anesthesia | 30 min, 150 min post-procedure |
| Castration | 30 min, 150 min, 300 min post-procedure |

**Fig 2.**
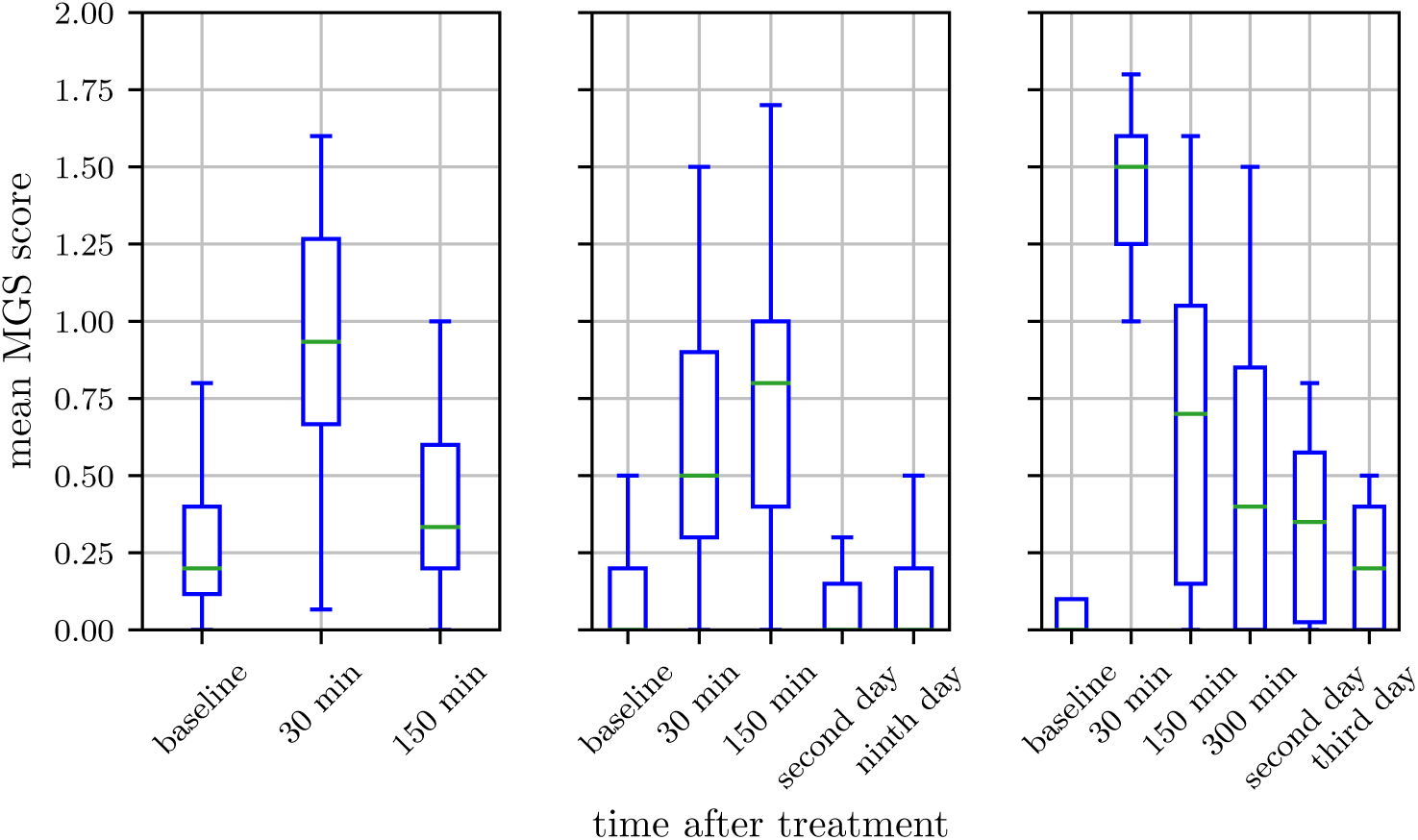
Box plots of the human evaluated Mouse Grimace Scale. Isoflurane anesthesia (IN, left), ketamine/xylazine anesthesia (KXN, middle) and castration (C, right). IN: Scores were obtained from 30 female and 31 male C57BL/6JRj mice. KXN: Scores were obtained from 29 female and 32 male C57BL/6JRj mice. C: Scores were obtained from 19 male C57BL/6JRj mice. Data represent the mean MGS scores averaged over three human scorers. The box represents the interquartile range (IQR), box edges are the 25th and 75th percentile. The whiskers represent values which are no greater than 1.5 × IQR. Outliers were excluded from the figure. This figure contains data from Hohlbaum et al. [29, 30].

#### Face detection

The first step of the evaluation pipeline is to distinguish between valid and invalid images. Valid images are those which contain a mouse face including at least one of the relevant features (eyes, ears, whiskers, nose, cheek) for facial expression analysis. Invalid images do not show a face or are of low quality (i.e., blurry). As not only the mouse but also parts of the observation cage are shown in the images, it was furthermore necessary to crop the relevant area of an image. This facilitates the learning of a mapping between image and label as a greater fraction of the input dimensions (pixels) of the model correlate with the target. Therefore, an automatic face detector was applied.

Sotocina et al. [6] created the “Rodent Face Finder” for white rats which was used in Tuttle et al. [36]. It combines two detectors, one for ears and one for eyes. Groups of detections are filtered according to heuristic expectations of a typical face (e.g., the ears must be above the eyes). Unfortunately, the trained model, provided by the authors of the “Rodent Face Finder”, led to a detection rate of zero, when applied to our data set. This is probably due to the different fur color of the mice in the two data sets. The training of a similar detector on our data set combined with the heuristics from Sotocina et al. resulted in a detection (face hypothesis) for 40% of all images while about 86% of all images are valid. We suppose that the “Rodent Face Finder” has a similarly low detection rate on white mice, but for the application on video sequences, as in Sotocina et al. [6] and Tuttle et al. [36], this is not a hindrance as a face needs only to be successfully detected in a fraction of the acquired frames for a correct classification. If a frame of the face is grabbed from a video every few seconds, the classification can be performed accurately. However, to achieve a bigger resulting training set for the facial expression recognition on our data set of still images, we decided to use a face detector with higher sensitivity and lower precision. As in Sotocina et al., we used a detector based on Boosted Cascades of Simple Features as introduced in Viola and Jones [37]. We used an implementation from the Computer Vision library OpenCV [38]. In contrast to the “Rodent Face Finder”, it was trained to detect the whole mouse face. Producing a higher false positive rate, this resulted in face hypotheses for about 80% of the images. An additional application of the eye and ear detector on the remaining 20% of images led to face hypotheses for 92% of the images. We applied these detectors to all 32576 images of 61 female and 65 male C57BL/6JRj mice and cropped the images based on the detected bounding boxes around the mouse faces. Afterwards, we manually sorted out false positives and images of low quality, which resulted in a remaining set of 18273 images (13352 from KXN, 2470 from C, and 2451 from IN; 60 female and 64 male C57BL/6JRj mice) (This is the resulting set we provide under www.scienceofintelligence.de/research/data/black-mice).

### Facial expression recognition

As in Tuttle et al. [36], we trained a binary classifier to distinguish between images labeled with “pain” or “no pain”. The binary classifier was a single layer fully connected neural network on top of a ResNet50 or InceptionV3 architecture. Note that in the study Tuttle et al. [36] only the InceptionV3 architecture was used. The weights of the ResNet50 and InceptionV3 networks were pre-trained on ImageNet [39] and frozen during training of the top layer. The models were chosen due to their current success. The InceptionV3 [40] model was tested on the ILSVRC 2012 [41] data set and achieved 21.2% top-1 error rate. The ResNet50 [42] achieved 20.1% top-1 error rate, which corresponds to the current state of the art in image-based object recognition. Additionally, we compared to a simpler architecture without pre-training (Fig 3). For training we used the Adam optimizer (parameters for Adam optimizer: learning rate 0.001, beta1 0.9, beta2 0.999, epsilon 1e-07, decay 0) to minimize the categorical cross entropy loss. All networks were implemented and trained using TensorFlow [43]. The output of the top layer of the network are the activations of two output neurons, one for each class. We classify an image as “pain” if the activation of the “pain” neuron is greater than the activation of the “no pain” neuron and vice versa. Furthermore, we interpret the resulting activation of the two output neurons as the confidence of the network for the two classes “pain” and “no pain”.

**Fig 3.**
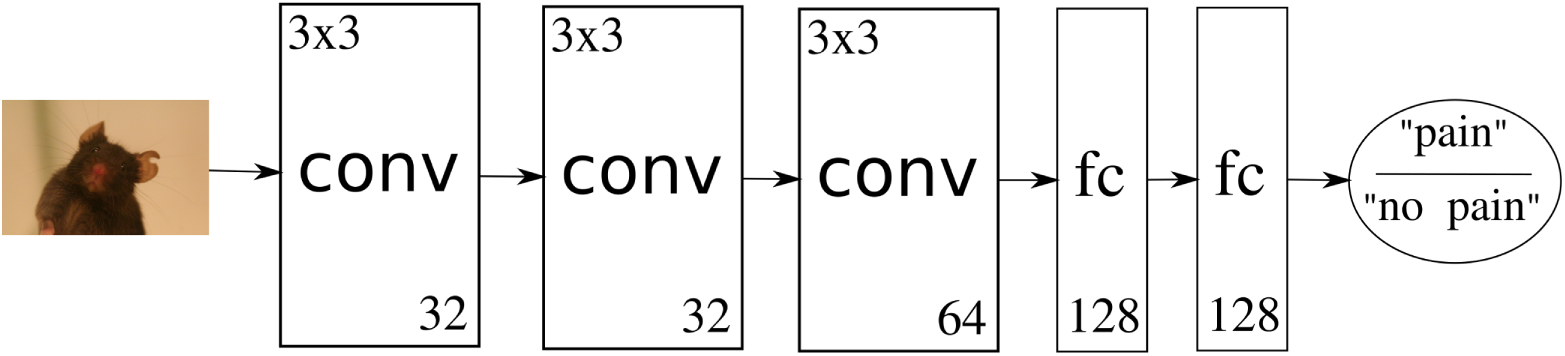
Own Network Architecture. The image is fed through three convolutional layers with filter size 3×3 and 32, 32 and 64 filters. Two fully connected layers with 128 neurons each follow. The two output neurons give a confidence in either judgment.

The network was trained with batch size 100 until it converged after 50 epochs using a new random permutation of the training set for each epoch. The data set was split in training and test set with no subject overlap for most experiments. In this context, the term subject stands for mouse. This strict separation ensures that no specific features of an individual are used for classification. It furthermore avoids the possibility of images with high similarity (taken at the same time of the same mouse) in training and test set. As 19 male mice which received ketamine/xylazine anesthesia or no treatment were reused for castration, some experiments have overlapping subjects. However, high similarity of images is ruled out here as images were acquired under different conditions (respective results are marked with superscript SO). To be able to train on the precomputed features and as the top layer did not overfit the data, we applied no data augmentation. The data sets were balanced by sub-sampling. All images were resized to 224 × 224 using bilinear transform.

## Results and Discussion

### ”Pain-No-Pain” Classification

First we evaluated the performance of the trained classifier, once on a combined set of all images and furthermore on the three subsets KXN, IN and C separately. We performed 10-fold cross-validation for the combined set and on the subsets KXN and IN and leave-one-animal-out-cross-validation for the subset C, which contains only 19 animals in total (Table 2a, Table 2b, Table 2c). The split into training and test set resulted in 112 animals for training and 12 animals for testing over all treatments, in 55 animals for training and 6 animals for testing for the KXN subset and in 57 animals for training and 6 animals for testing for the IN subset.

**Table 2.** Cross-validation results. Data are given as mean percent ± standard deviation. IN: isoflurane anesthesia; KXN: ketamine/xylazine anesthesia; C: castration; TPR: true positive rate (sensitivity); TNR: true negative rate (specificity)

|  | accuracy | TPR | TNR |
| --- | --- | --- | --- |
| All | 88.5 ± 2.2 | 85.8 ± 4.4 | 90.8 ± 2.6 |
| KXN | 93.4 ± 1.9 | 93.4 ± 4.2 | 92.4 ± 4.4 |
| C | 89.5 ± 3.6 | 89.6 ± 9.0 | 89.2 ± 5.7 |
| IN | 75.6 ± 4.8 | 74.4 ± 7.9 | 76.6 ± 6.3 |

(b) Inception.
|  | accuracy | TPR | TNR |
| --- | --- | --- | --- |
| All | 83.8 ± 3.1 | 82.1 ± 4.7 | 85.1 ± 5.7 |
| KXN | 90.0 ± 2.4 | 86.3 ± 6.1 | 92.1 ± 3.1 |
| C | 78.0 ± 8.9 | 74.8 ± 17.1 | 81.2 ± 10.0 |
| IN | 72.4 ± 4.7 | 67.4 ± 11.6 | 75.4 ± 9.1 |

(c) Own architecture.
|  | accuracy | TPR | TNR |
| --- | --- | --- | --- |
| All | 82.9 ± 4.2 | 81.3 ± 7.7 | 83.3 ± 5.0 |
| KXN | 88.1 ± 2.5 | 87.1 ± 4.7 | 90.4 ± 4.5 |
| C | 83.5 ± 3.0 | 82.3 ± 5.7 | 84.3 ± 5.1 |
| IN | 56.5 ± 4.7 | 62.8 ± 12.2 | 55.8 ± 13.8 |

(d) ResNet. Averaging over multiple images.
|  | accuracy | TPR | TNR |
| --- | --- | --- | --- |
| All | 92.5 ± 2.4 | 86.3 ± 7.2 | 95.8 ± 2.0 |
| KXN | 98.9 ± 1.9 | 97.2 ± 4.7 | 100.0 ± 0.0 |
| C | 89.8 ± 5.5 | 98.1 ± 7.6 | 85.3 ± 9.2 |
| IN | 90.1 ± 5.7 | 93.5 ± 10.0 | 89.0 ± 9.5 |

Although the InceptionV3 architecture was used in previous works [36], we achieved highest values for accuracy (= *t p*+*tn/t p*+*tn*+ *f p*+ *f n*), sensitivity (= *t p/t p*+ *f n*), and specificity (= *tn/tn*+ *f p*) for the combined data sets on the ResNet architecture. However, if we evaluated the data sets separately, performance differed for KXN, C, and IN. While performance improved for KXN as well as C, it deteriorated for IN. To understand why results were lower for IN, we have to consider the design of the study from which images of the IN subset were obtained. Mice of this study received inhalation anesthesia with isoflurane, which caused statistically significant changes in the facial expression for a relatively short period only [29]. Therefore, images taken 30 min post-anesthesia were considered to display a “pain” face and images generated at baseline or 150 min post-anesthesia were labeled with “no pain”. Since our previous study did not reveal any statistically difference in the facial expressions according to the MGS between the time points baseline and 150 min post-anesthesia [29], MGS scores were still slightly increased in some animals at 150 min post-anesthesia. As a consequence, the binary classification led to a pool of “no pain” images with a high range of intensities (Fig 2, S3 Fig). The “no pain” and “pain” classes include MGS scores of 0.20 (median; interquartile range: 0.28) and 0.67 (median; interquartile range: 0.73), respectively. The resulting smaller margin in feature space for this treatment would probably require a classifier with higher complexity compared to the other treatments.

As Tuttle et al. [36] showed that results can be improved by using multiple frames from video data for classification, we too averaged the network confidence over all images that were taken at the same point in time (i.e., have same time label) of the same animal and then classified the images based on the average confidence (Table 2d). As the images were taken from different perspectives the additional information here is higher than in the use of video data where the frames potentially contain a lot of redundant information. The results show increased performance for all subsets, especially IN. For the subset KXN, specificity was 1.000 ± 0.000 (mean ± standard deviation). This high result may be explained by a clear decision boundary between the two classes in the KXN subset. Injection anesthesia is known to intensively impair the general condition of a mouse and to significantly affect its facial expression for a longer period (i.e., up to at least 150 min) [30]. Therefore, images generated 30 min as well as 150 min post-anesthesia were labeled with “pain” and images of the remaining time points were assigned the label “no pain”. In contrast to the subset IN, the range of intensities of facial expressions in the “no pain” class is smaller in the subset KXN (Fig 2, S3 Fig). MGS scores of the “no pain” class (median: 0.00, interquartile range: 0.2) and “pain” class (median: 0.60, interquartile range: 0.60) overlap to lesser extent than seen for IN, which may have contributed to a clearer decision boundary between the two classes.

The performance of the algorithms in binary classification of “pain” versus “no pain” images cannot be directly compared to the human performance in our data set because we used the MGS scores obtained from humans as ground truth. Langford et al. [5] reported an accuracy of 97% and 81% for experienced and inexperienced human scorers, respectively, when high resolution images (1,920 × 1,080 pixels) were used. A lower accuracy of 72% was found for inexperienced humans who scored low resolution images (640 × 480 pixels) [5]. This underlines the importance of both image quality and experience in the use of the MGS for the application of this method by humans. Images of our data set originally had higher resolution (3456 × 2304 pixels), but were resized to 224 × 224 for the present study in order to reduce input dimensionality of the learner. All tested architectures showed higher accuracy on the combined data set of all images (i.e., including IN, KXN, and C) when compared to inexperience humans using low or high resolution images. Similar success to experienced human scorers was only achieved by ResNet for the subset KXN when a classification was based on all available images of an animal at a certain point in time (Table 2d).

### Network confidence over time

In Fig. 4, Fig. 5, and Fig. 6, we present the network confidence values for the “pain” class of ResNet architecture and the human evaluated MGS scores over time for the subset IN, KXN, and C, respectively. In Tuttle et al. [36], the correlation analysis between the confidence of the classifier and the MGS score suggests, that an estimate of the expression intensity can potentially be inferred without the necessity of a regression model trained on MGS score labels. Since the intersection of images scored on the MGS and tested images in our data set is too small, a correlation analysis like in Tuttle et al. could not be carried out.

**Fig 4.**
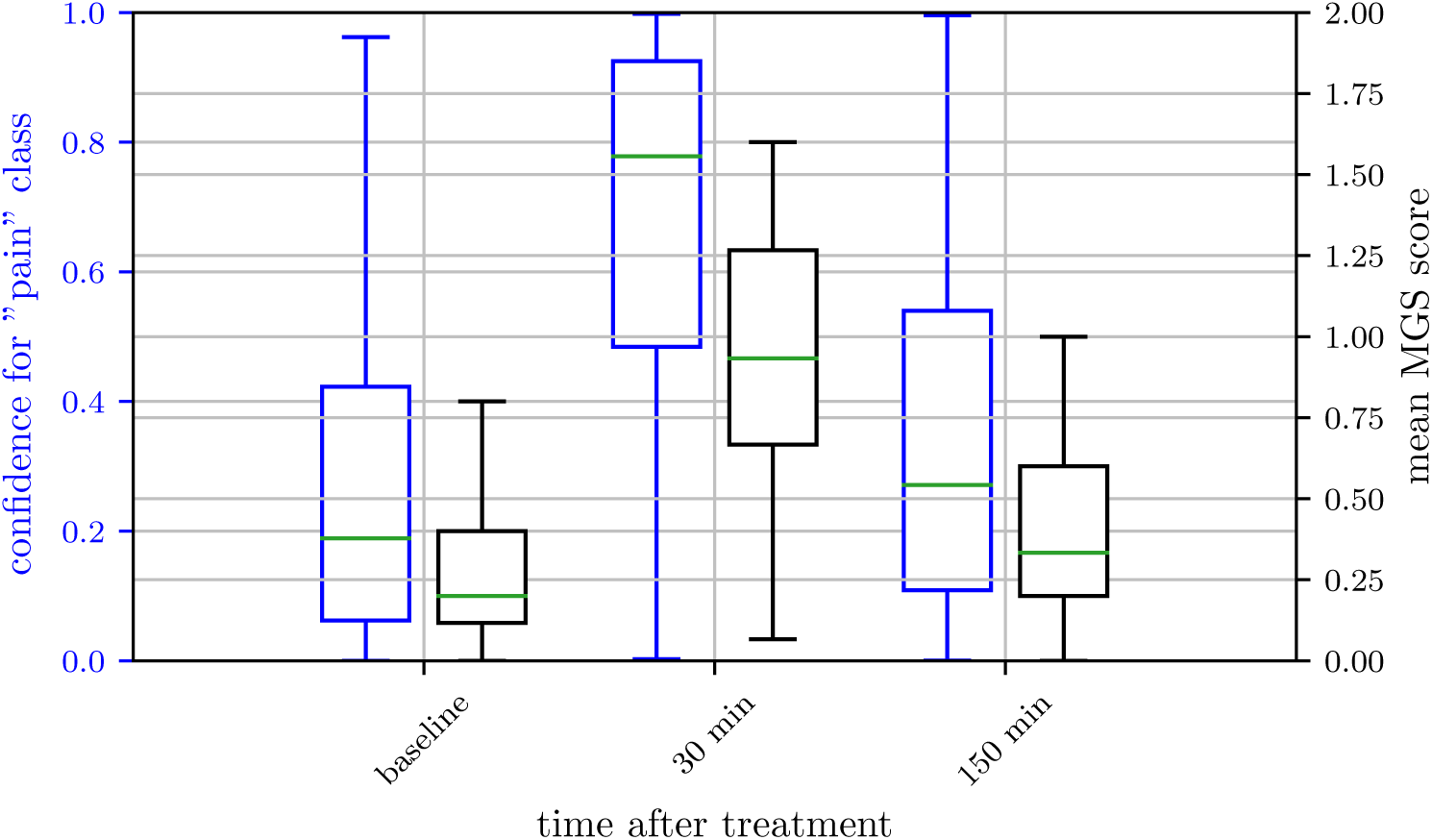
Network confidence over time for isoflurane anesthesia. plots of human labeled Mouse Grimace Scale (MGS) scores (grey) and confidence for “pain” class of ResNet architecture (blue) for isoflurane anesthesia (IN). Scores were obtained from 33 female and 32 male C57BL/6JRj mice. MGS data represent the mean MGS scores averaged over three human scorers. The box represents the interquartile range (IQR), box edges are the 25th and 75th percentile. The whiskers represent values which are no greater than 1.5 × IQR. Outliers were excluded from the figure. This figure contains data from Hohlbaum et al. [29].

**Fig 5.**
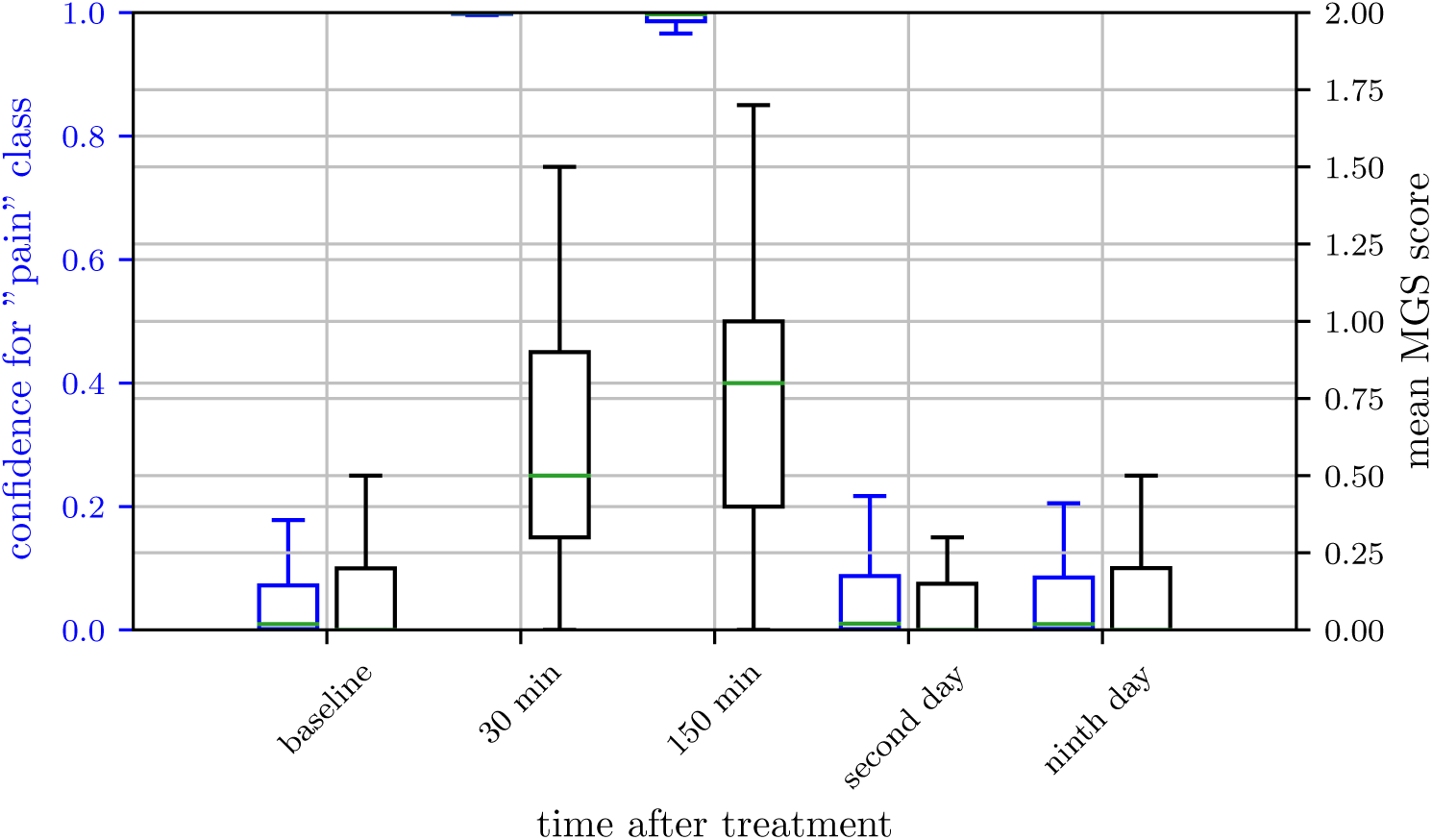
Network confidence over time for ketamine/xylazine anesthesia. Box plots of human labeled Mouse Grimace Scale (MGS) score (grey) and confidence for “pain” class of ResNet architecture (blue) for ketamine/xylazine anesthesia (KXN). Scores were obtained from 28 female and 30 male C57BL/6JRj mice. MGS data represent mean MGS scores averaged over four human scorers. The box represents the interquartile range (IQR), box edges are the 25th and 75th percentile. The whiskers represent values which are no greater than 1.5 × IQR. Outliers were excluded from the figure. This figure contains data from Hohlbaum et al. [30].

**Fig 6.**
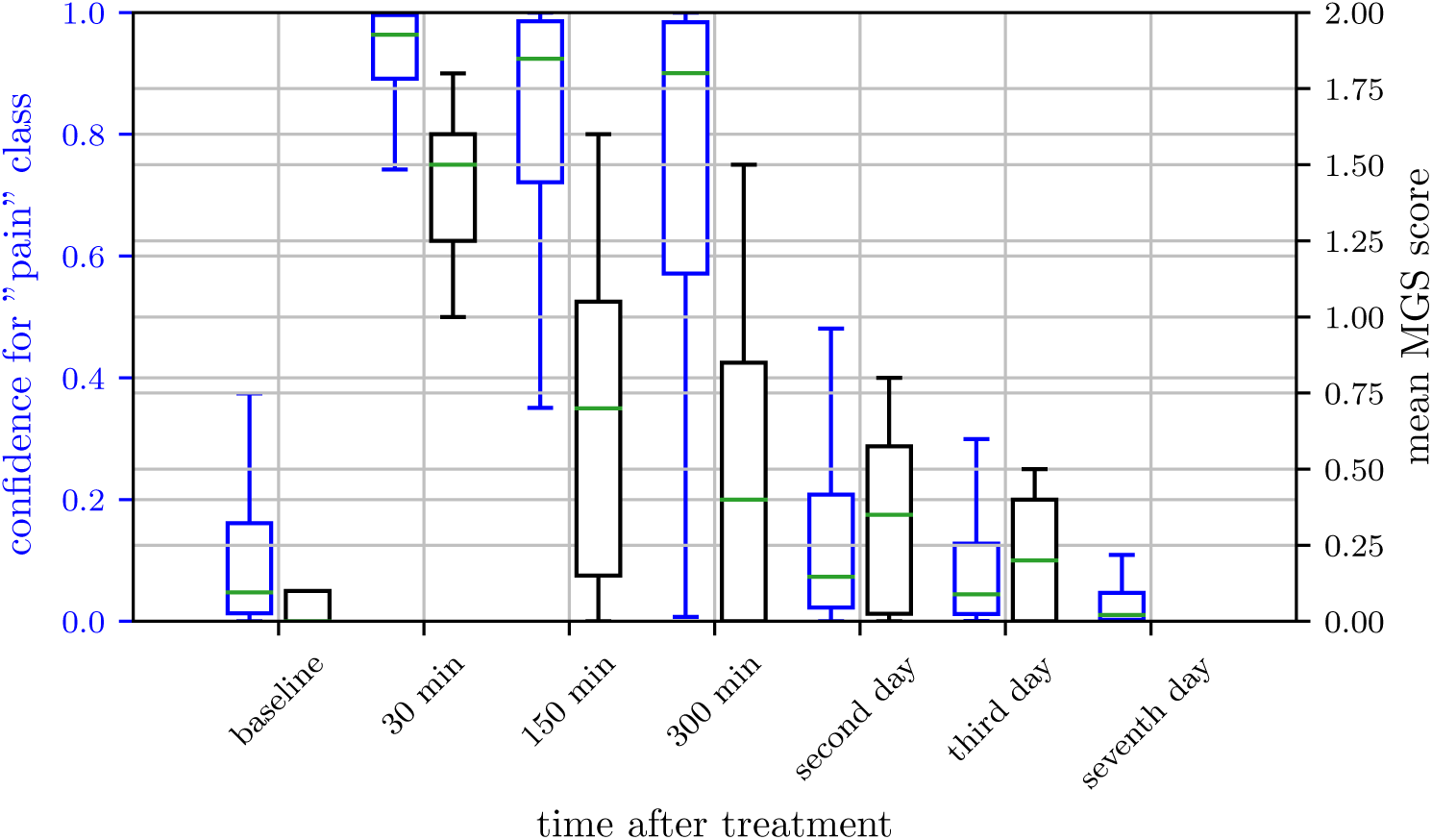
Network confidence over time for castration. Box plots of human labeled Mouse Grimace Scale (MGS) score (grey) and confidence for “pain” class of ResNet architecture (blue) for castration (C). Scores were obtained from 19 male C57BL/6JRj mice. MGS data represent the mean MGS scores averaged over two human scorers. The box represents the interquartile range (IQR), box edges are the 25th and 75th percentile. The whiskers represent values which are no greater than 1.5 × IQR. Outliers were excluded from the figure.

However, in general, the data suggests that the confidence for the “pain” class was higher for images with high MGS scores. This can be clearly seen for 30 min and 150 min post-anesthesia with ketamine/xylazine combination (Fig 5) or 30 min post-castration (Fig 6). The other way around, a very low confidence for the “pain” class was found for images with low MGS scores, indicating a “no pain” state. In intermediate cases however, while the network output still tends to follow the human evaluated MGS score, the deviation of network confidence and MGS score increases (e.g. 150 min and 300 min post-castration).

Regarding inhalation anesthesia with isoflurane, the confidence for the “pain” class reflects the difficult decision boundary between the two classes “pain” and “no pain” in this data subset (Fig 4), as discussed above. In brief, both classes “pain” and “no pain” contain images within a relatively big, shared range of the MGS scores, what makes it more difficult for the algorithm to distinguish between the two classes.

### Cross-Treatment Evaluation

To figure out if there is a universal “pain” facial expression independent of treatment, we performed cross-treatment analysis of our algorithms. In Fig 7 the accuracies of the ResNet50 and InceptionV3 architectures are presented. Although their absolute performance is different, the relative performance for various combinations of training and test data sets seems to be similar. For a fair comparison these results are based on a subset of the data available for KXN with the same size as the C and IN sets. It was noticeable that the performance of both algorithms decreased in nearly all cases when they were trained and tested on mouse images of different treatments. To explain this, we developed two hypotheses regarding the dimensionality of the expression space. First, the learner may learn different decision boundaries for different distributions of facial expression intensities in different treatments and therefore performs poorly on the other subsets. This might be counteracted by a better design of the label space. Secondly, the expression space is inherently multidimensional and the learned interpretation cannot be transferred between different treatments. This is harder to overcome but would potentially allow to make higher dimensional estimates for the state of the animal. Treatments underlying our data set, i.e., inhalation anesthesia, injection anesthesia, and castration, may induce different facial expressions. This hypothesis is supported by the effects of these treatments on the general condition of a mouse. In general, surgery including inhalation anesthesia and analgesic treatment, depending on analgesia management, causes a higher impairment of well-being than inhalation anesthesia only, which can be assessed by behavioral parameters, for instance nest building [44]. Depending on the treatment, a mouse is exposed to different stimuli and experiences different states. Post-surgical pain accompanies castration, whereas anesthesia is unlikely to induce pain. However, the injection with the combination of ketamine and xylazine can damage the tissue at the injection site, which may be associated with a painful sensation [45], but we assume the degree of pain to be much lower when compared to post-surgical pain. Anesthesia induces post-anesthetic distress with a variety of causes. Isoflurane has a pungent odor [46] and induces irritant effects in the airways by activating nociceptive ion channels [47, 48]. In humans, the inhalation of isoflurane causes coughing and subjective sensations of burning as well as irritation [49]. In addition, if 100% oxygen is used as carrier gas, the inhalation gas is very dry and can impair the function of the respiratory mucosa [50]. When anesthesia is induced, distress of a mouse can additionally increase by fixation and injection stress or by exposure to the (irritant) volatile anesthetic. In the latter case, aversion towards this inhalant agent elevates with repeated exposure [51]. Distress a mouse experienced during the induction phase may influence its well-being after anesthesia as well. When mice recover from anesthesia, they can suffer from post-anesthetic nausea [52]. Moreover, in humans emergence delirium can occur during the recovery period and hallucinogenic effects were reported for the use of ketamine [46, 53]. We also have to consider different pharmacological effects of anesthetics on the facial expressions of the mice with longer lasting effects following injection anesthesia due to the pharmacokinetic properties of ketamine and xylazine. In contrast to isoflurane, ketamine and xylazine are subject of an intensive liver metabolism [54–57], which results in longer recovery periods. Ketamine increases the muscle tone, whereas the combination of ketamine and xylazine causes muscle relaxation [46]. All in all, inhalation anesthesia, injection anesthesia, and castration produce different affectional states in a mouse. Against the background that the weight of the five facial action units varies between different states like illness and pain [5, 58], our data suggests that the procedures we investigated in the present study induce different facial expressions. This may explain the reduction in performance when the algorithms were trained and tested on mouse images of different treatments and is a disadvantage for the pure binary classification of “pain” and “no pain”.

**Fig 7.**
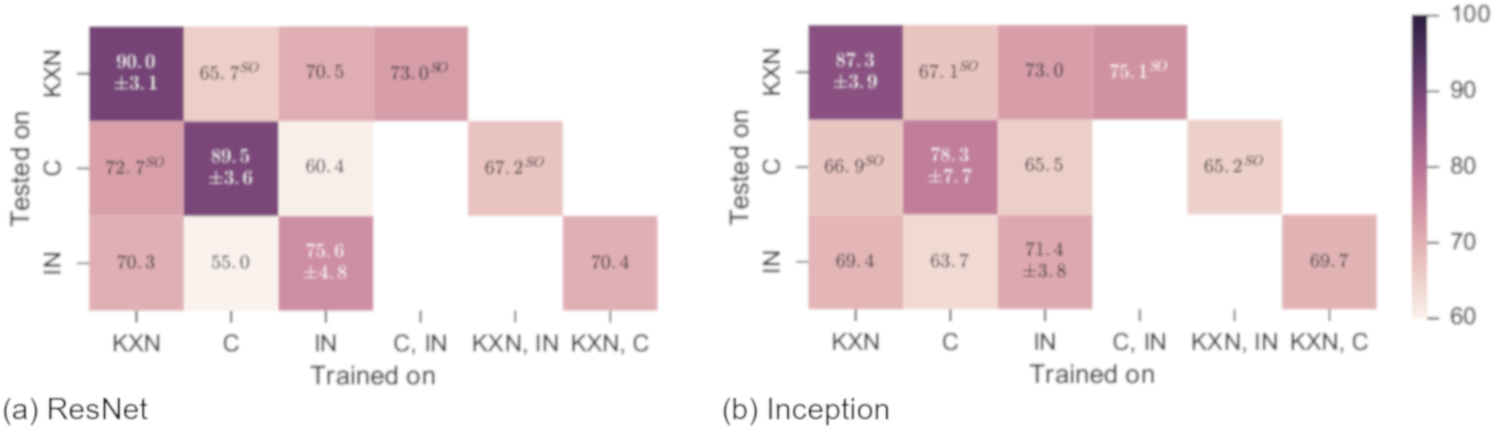
Performance for different combination of training and test datasets after 50 epoch of training. These results are based on a subset of the data available for KXN. The subset has the same size as the K and IN sets and allows a fair comparison of the values. Data are given as mean accuracy (± standard deviation) in %. IN: isoflurane anesthesia; KXN: ketamine/xylazine anesthesia; C: castration; SO: subject overlap (the term subject is used as a synonym for mouse).

### Feature importance visualization

To ensure that the predictions of the networks are based on the facial expressions of the mice and not on other high level (e.g., food pellets, ear punches used as markers) or low level (e.g., illumination) features that possibly correlate with the target (”pain”/”no pain” state) and to get insights into which features the neural network uses for classification, we performed a decomposition analysis. Methods as the deep Taylor decomposition [59–61], which we used in this work, propagate the activations of the neurons backwards through the network and decompose the prediction into contributions of single pixels. These contributions can be visualized as heat maps to explain the decision of a network regarding a certain image. From Fig. 8 we clearly see, that the decision is made using mostly features related to the mouse itself and not to the background. The ears and especially the outline of the ears, the eyes and the area around the eyes, the nose tip and the area around the nose (i.e., probably the whisker pad), as well as whiskers or the space between the whiskers are contributing to the decision making by the neural network. Thus, the heatmaps confirm the importance of at least some facial action units of the MGS [5], especially orbital tightening, ear position and whisker change.

Additionally, further facial features such as the nose tip and the whisker pad appear to have a significant impact on the decision making process, particularly in mouse images labeled with “no pain” (S1 Fig). To understand why the nose tip seems to play an important role in the “no pain” state, but less in the “pain” state, the color of the nose and its position is of special interest. While the nose points forwards or is slightly elevated in the “no pain” state in general, the head rather is dropped and the nose tip points downwards after anesthesia or surgery. The nose tip is colored (pale) pink in mice in good general condition. If mice recover from anesthesia or surgery, circulation can be affected in the early post-anesthetic period, hence the color of the nose may turn paler. Overall, the nose tip may play a more important role in the “no pain” state because it is clearly visible and the color is very prominent. S1 Fig and Fig. 8 (bottom row) reveal that, except from the nose tip, the area around the nose, probably the whisker pad, can also be critical for the decision making process. The importance of this feature may be traced back to the muscles associated with the vibrissal follicles [62], which cause a change in whisker position. A natural downward curve of the whiskers is found in the “no pain” state, whereas whiskers stiffen and are pulled back or forward, they may also clump together in the “pain” state. Besides the whisker position, the activity of these muscles may influence the appearance of the whisker pad, which would explain the usefulness of this feature for the decision making. Another relevant feature we detected by visualization was the space between the whiskers, which is obviously influenced by the position of the whiskers and may have the same meaning accordingly (S2 Fig). However, the network uses not only pixels of the mouse face but also pixels of the body if visible, such as the outline of the back or in some cases the cervical/thoracic area (Fig 8). The outline of the back is altered if a mouse shows a hunched posture or piloerection (S2 Fig), which can accompany distress and pain [63]. The cervical/thoracic area is only entirely visible if the animal sits upright, which can be used as an additional clue for the state of the mouse, i.e., the head of a mouse lies flat on the ground in most images taken in the early post-anesthetic period following injection anesthesia with ketamine/xylazine combination. Moreover, in images from this period particular facial features are very distinctive, e.g., the ear position. Probably due to pharmacological effects of the injection anesthetics, the ears are dropped and the space between the ears widens. In some cases, decomposition analysis even suggests that the decision for the classification is mainly based on the ears (see bottom row in S1 Fig).

**Fig 8.**
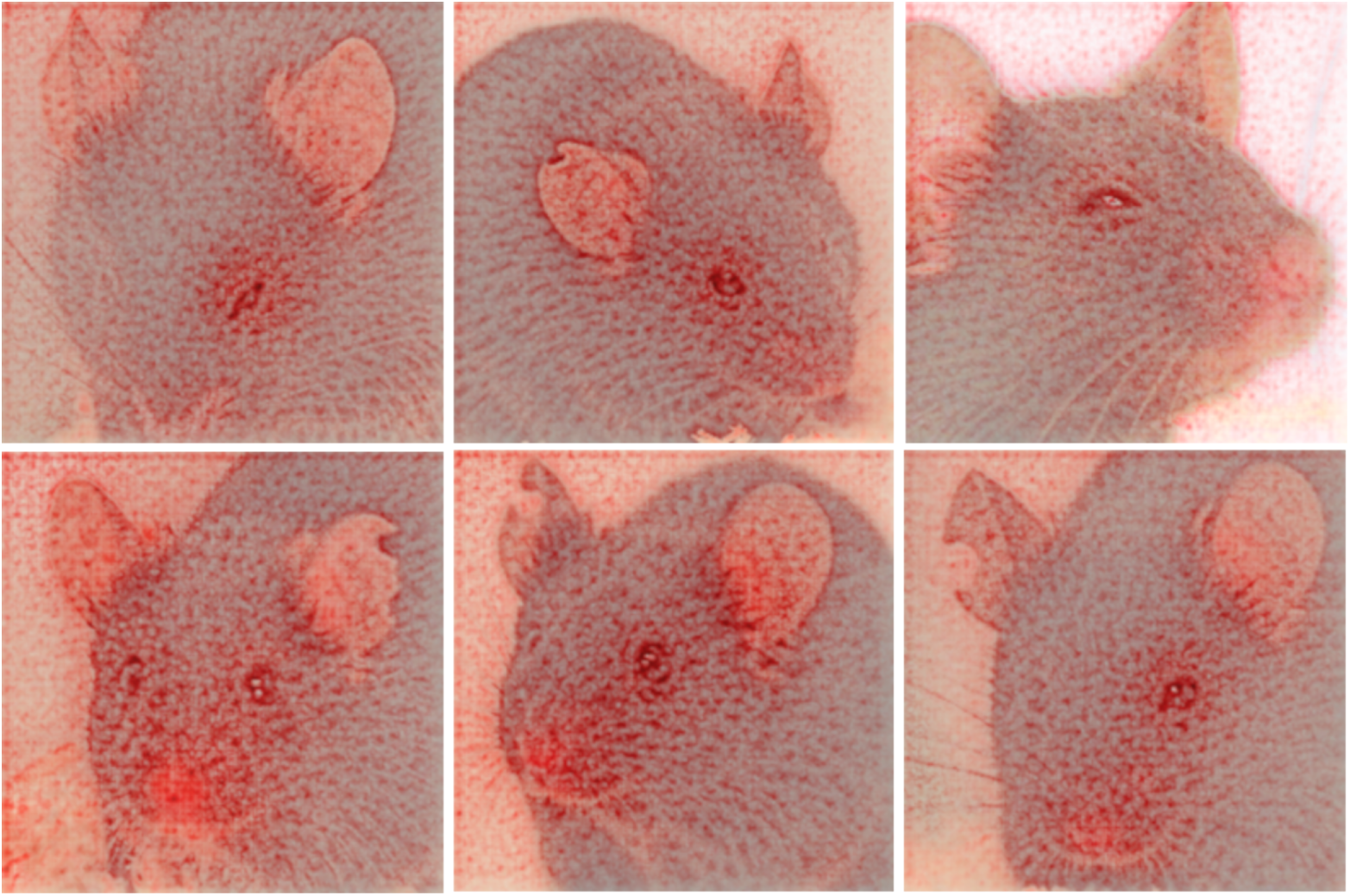
Visualization of the decision finding process using deep Taylor decomposition. Castration (left), ketamine/xylazine anesthesia (middle), and isoflurane anesthesia (right). Mice correctly classified with “pain” in top row, mice correctly classified with “no pain” in bottom row. Red color indicates that a pixel contributes to the decision.

However, in order to determine which of the various facial and body features contributed most to the decision making progress of the deep neural network, future work is needed. We hope that we will be able to localize facial elements and body parts in the images. This would enable us to quantitatively analyze the weighting of the used features and allow to compare features of the algorithm with the facial action units of the MGS. It would furthermore enable us to investigate whether the decision for models trained on different procedures (i.e., IN, KXN, C) is based on different features. These information may lead to a better understanding of the high confidence of the network in cases such as 30 min and 150 min post-anesthesia with the combination of ketamine/xylazine.

## Conclusion

We developed a semi-automated pipeline to automatically recognize post-anesthetic or post-surgical distress and/or pain of a mouse from its face using a deep learning neural network framework. For the first time, this approach was pursued involving images of black-furred laboratory mice moving freely in their cages. Depending on the treatment of the mice, an accuracy of up to 99% was reached for the binary classification (”pain”/”no pain”). A model trained on a particular treatment performs worse if it is tested to another treatment, suggesting that different treatments produce divergent facial expressions. Mainly eyes, ears, and whiskers, but furthermore in some cases additional facial (nose tip, whisker pad) as well as body (outline of the back, cervical/thoracic area) features seem to contribute to binary classification in some cases. The findings of our study promote the development of a prototype tool for monitoring well-being of laboratory mice in experimental settings and their home cages.

### Outlook

The long term goal is to devise a smart-surveilled environment for laboratory mice. The proposed approach will be the foundation for a “smart mouse cage”, i.e., an integrated system with around-the-clock video monitoring of laboratory mice and alerting staff, if the well-being of an animal is impaired. Another possible application will be a computer program or smart phone app, which supports the experimenter in assessing MGS scores in real time. The envisaged app could be used for online analysis of video stream with alert generation in case of a deviance appearance, i.e., a pain face. We believe that our approach for automated surveillance of the well-being state would also be useful for other animal species. However, one of the crucial steps for pipeline automation is a proper face finder. The one currently used would need to be replaced in order to not require manual rejection of false positives (e.g. with Mathis et al. [64]). The use of video data instead of still images could also lead to a fully automated face detection as face hypotheses with low confidence scores could be rejected. Another interesting future direction of research would be an assessment of relative importance of features and its comparison to the MGS scoring scheme, which may reveal new facial features beyond the MGS. Besides negative well-being, facial expressions may also represent positive well-being. While facial indicators of positive emotions in mice have not been reported yet, it might be achieved with the help of an automated pipeline.

## Acknowledgments

The authors would like to thank Doris Ciuraj, Sabine Jacobs, Melanie Humpenöder, and Giuliano Corte for assisting in MGS scoring. We are grateful to Oliver Leonhardt who developed our MGS scoring database. The study was funded by the Deutsche Forschungsgemeinschaft (DFG, German Research Foundation: www.dfg.de) under Germany’s Excellence Strategy – EXC 2002 “Science of Intelligence” – project number 390523135 (www.scienceofintelligence.de). Mouse Grimace Scale scores were obtained from a project of the Berlin-Brandenburg research platform BB3R (www.bb3r.de) that was funded by the German Federal Ministry of Education and Research (grant number: 031A262A) (www.bmbf.de/en/index.html).

This work’s data is released open access and can be found under www.scienceofintelligence.de/research/data/black-mice.

## Supporting information

**S1 Fig.**
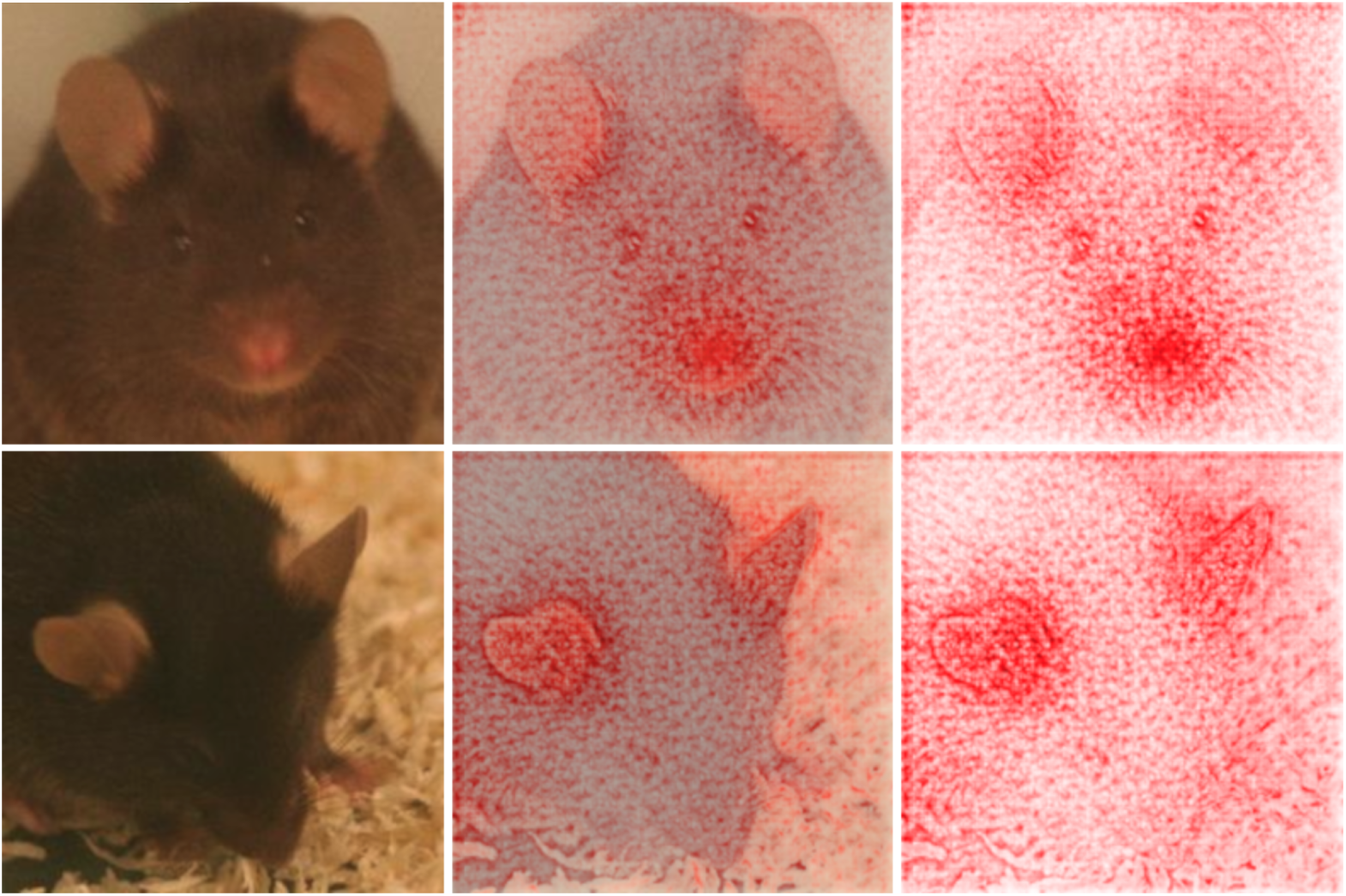
Contribution of nose, whisker pad, and ears to the decision making. Visualization of the decision finding process using deep Taylor decomposition for images generated 30 min (top row) and 2 days (bottom row) after ketamine/xylazine anesthesia. Original image (left), original image combined with heat map (middle), heat map (right). Red color indicates that a pixel contributes to the decision. Top row: The mice was correctly classified as “no pain” with a confidence of 96,2%. In particular the nose and the whisker pad seem to contribute to the decision. Bottom row: The mice was correctly classified as “pain” with a confidence of 100,0%. The decision appears to be mainly based on the ears.).

**S2 Fig.**
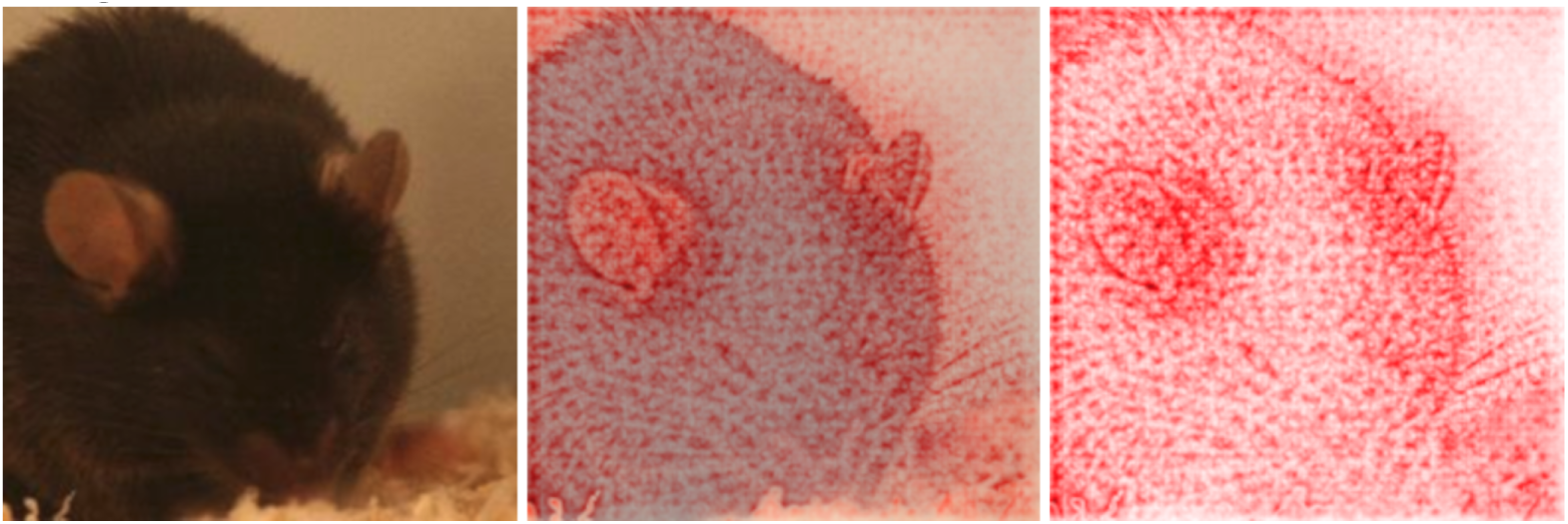
Contribution of piloerection and space between whiskers to the decision making. Visualization of the decision finding process using deep Taylor decomposition for an image generated 150 min after ketamine/xylazine anesthesia. Original image (left), original image combined with heat map (middle), heat map (right). Red color indicates that a pixel contributes to the decision. Piloerection and the space between the whiskers seem to play a role in the decision making progress. The mice was correctly classified as “pain” with a confidence of 99,6%.

**S3 Fig.**
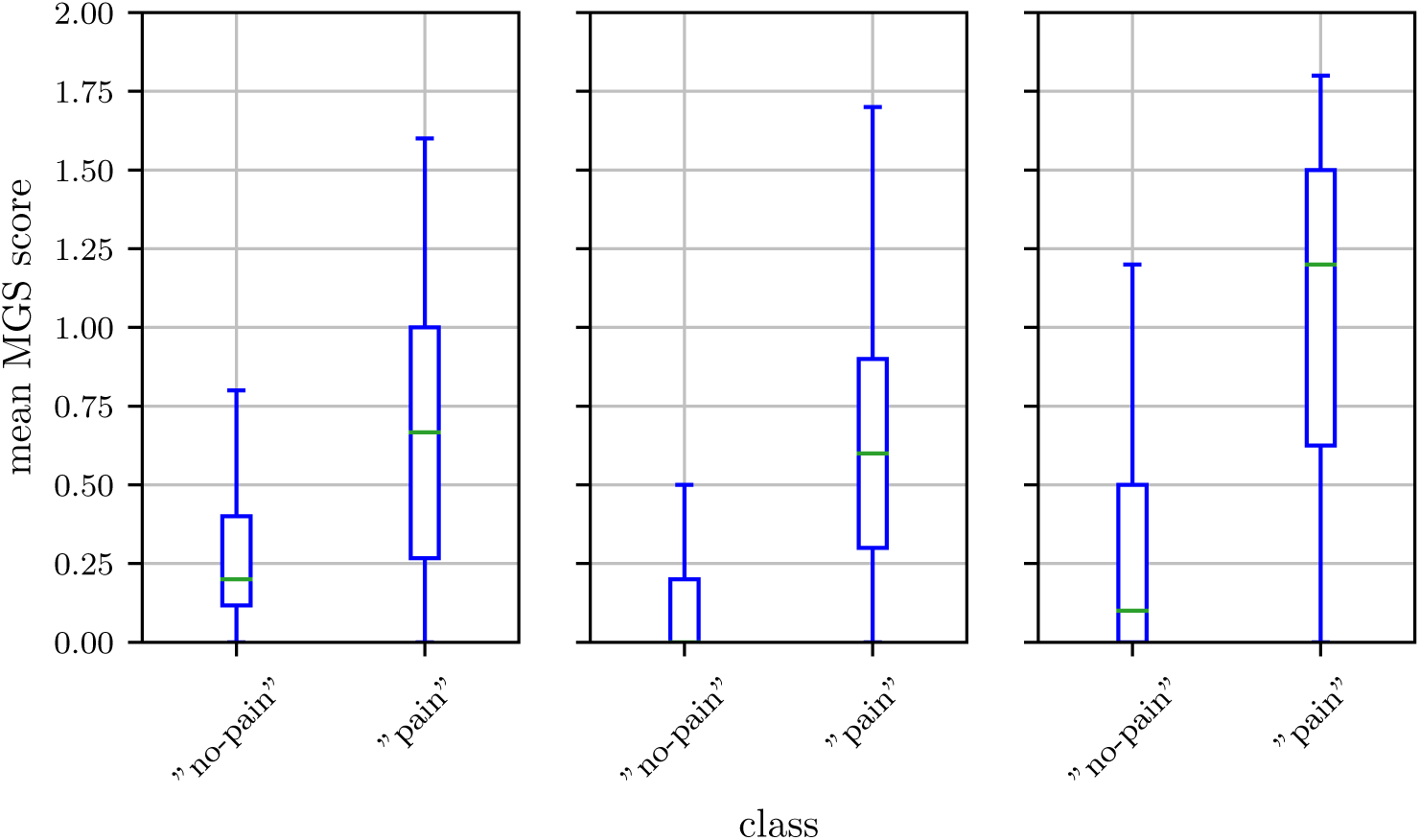
Ground truth distribution of the “pain” and “no pain” class. Isoflurane anesthesia (IN, left), ketamine/xylazine anesthesia (KXN, middle), and castration (C, right). The box represents the interquartile range (IQR), box edges are the 25th and 75th percentile. The whiskers represent values which are no greater than 1.5 × IQR. Outliers were excluded from the figure.

